# Repressing ABCB7 Potentiates Cisplatin Response in Pediatric Group 3 Medulloblastoma by Triggering Ferroptosis

**DOI:** 10.1101/2022.01.24.477587

**Authors:** Ranjana K. Kanchan, Naveenkumar Perumal, Parvez Khan, David Doss, Prakadeeshwari Gopalakrishnan, Ramakanth Chirravuri Venkata, Ishwor Thapa, Raghupathy Vengoji, Jyoti B. Kaushal, Jawed A. Siddiqui, Mohd Wasim Nasser, Surinder K. Batra, Sidharth Mahapatra

## Abstract

Medulloblastomas (MB) are the most common malignant pediatric brain tumor and a leading cause of childhood mortality. Aggressive tumors belonging to group 3 (G3MB) are distinguished by a marked reduction in programmed cell death by ferroptosis. These aggressive tumors also enrich iron transport and glutathione metabolism. A highly enriched pathway in these tumors is iron-sulfur (Fe-S) cluster binding corresponding to significant upregulation of ABCB7, a poor prognostic feature linked with accelerated mortality. This study elucidated whether repressing ABCB7 activates ferroptosis to mitigate G3MB aggressiveness and whether this pathway is pharmacologically targetable. *In silico* and *in vitro* analyses confirmed upregulation of ABCB7 and GPX4, the central regulator of ferroptosis, in G3MB cell lines and tumors. Repressing ABCB7 (miR-1253^OE^, siABCB7, sh-ABCB7) induced iron overload, elicited oxidative stress, and triggered lipid peroxidation, leading to an abrogation of medullosphere formation and cell death by ferroptosis. Intriguingly, fractionation studies revealed that ABCB7 repression abrogated GPX4 expression, most likely by GSH depletion. Repressing ABCB7 induced mitochondrial dysfunction and reduced oxidative phosphorylation. Cisplatin, a chemotherapeutic mainstay of G3MB, induces cell death by DNA crosslinking. In ABCB7 repressed cell lines, the IC_50_ of cisplatin was halved, resulting in augmented oxidative stress and lipid peroxidation, culminating in a higher index of ferroptosis. Artesunate, an anti-malarial drug capable of triggering ferroptosis, was shown to synergize with cisplatin, reducing tumor burden and significantly prolonging survival. Taken together, the current study illustrates how targeting iron transport can augment ferroptosis in G3MBs. It further identifies an FDA-approved drug capable of recapitulating these effects and potentiating mainstay chemotherapy.

## Introduction

Of the primitive neuro-ectodermal tumors (PNET) afflicting the posterior fossa, medulloblastoma (MB) is the most common malignant tumor of childhood, accounting for 85% of PNETs and 20% of all posterior fossa tumors.^1–3^ Based on high-throughput gene profiling studies, MBs are stratified into four major molecular subgroups: WNT, SHH, group 3 and group 4.^4^ Among these, patients with group 3 tumors (G3MB) suffer the worst prognosis (5-year overall survival <50%) due to a highly aggressive phenotype punctuated by c-Myc amplification, haploinsufficiency of chromosome 17p, presence of metastatic lesions at diagnosis, and high rates of recurrence.^4–10^ Recurrence can further reduce 5-year overall survival to <10% and is partially fueled by the inability of young patients to tolerate effective chemo-and/or radiation therapy.^4,11–13^ For instance, systemic toxicity is an oft-cited justification for limitations in cisplatin dosing, especially for group 3 disease, leading to <20% of treated patients receiving an effective dose.^14–16^ These facts underscore the urgent need to develop new strategies for high-risk disease.

Recently, combining microRNAs with chemotherapy has garnered support in enhancing therapeutic efficacy via targeting critical regulators of DNA damage repair (DDR), apoptosis, or cell cycle.^17^ In breast cancer, for example, miR-9 and miR-218 improved cisplatin responsiveness by targeting BRCA1 and impeding DNA damage repair.^18,19^ The miR-15 family sensitized cisplatin-resistant cancer cells to apoptosis by targeting the G_2_/M cell cycle checkpoint.^20^ In gastric and lung cancer cell lines, the miRNA cluster miR-200bc/429 sensitized resistant cell lines to both vincristine and cisplatin by targeting BCL2 and XIAP.^21^ In a gemcitabine-resistant pancreatic xenograft model, miR-205 delivered with gemcitabine in conjugated micelles resulted in significant tumor growth inhibition.^22^ Most notably, miR-34a replacement therapy (MRX34) has reached a phase 1 clinical trial (NCT01829971) in patients with primary unresectable or metastatic liver cancer.^23^

Group 3 tumors have amongst the highest frequency of cytogenetic aberrations targeting the short arm of chromosome 17 compared to the other subgroups, with high incidence reported on locus 17p13.3.^24–27^ We recently revealed strong tumor suppressive properties for miR-212 and miR-1253, which reside on locus 17p13.3.^27,28^ We showed that miR-1253 targets the cell cycle checkpoint protein, CDK6, and CD276 (B7-H3), an immune checkpoint molecule implicated in tumor aggressiveness. ^27,29–31^ Aside from reducing tumor cell viability, migration/invasion, and colony formation, miR-1253 arrested cells at the G_0_/G_1_ phase, activated apoptotic pathways, and triggered oxidative stress.^27^ Numerous studies have recapitulated these strong tumor suppressive properties in various cancers, including non-small cell lung carcinoma^32^, osteosarcoma^33^, pancreatic cancer^34^, male breast cancer^35^, and colon cancer^36^.

Amongst the putative targets of miR-1253 are the ATP-binding cassette (ABC) transporters that confer multi-drug resistance to various tumors, including MB.^37,38^ In metastatic breast cancer, for example, miR-1253 was shown to inhibit the drug efflux pump, ABCB1, thereby potentiating the cytotoxicity of doxorubicin.^39^ However, this particular ABC transporter is not amongst the deregulated ABC transporters reported in G3MB.^40^ Another putative target of miR-1253 is ABCB7, an iron transporter residing on the inner mitochondrial membrane involved in iron homeostasis and Fe-S cluster biogenesis.^41^ Deregulation of ABCB7 has been shown to help cancer cells withstand apoptosis and ferroptosis.^42^ Ferroptosis is a form of cell death triggered by iron-mediated oxidative stress leading to lethal lipid peroxidation. This process is tightly controlled by glutathione peroxidase 4 (GPX4), whose primary substrate is the anti-oxidant, glutathione (GSH).^43–45^ Recent studies have identified ferroptosis mitigation in tumor progression and drug resistance.^46,47^

Of the standard chemotherapeutic agents used in the treatment of MB, cisplatin, a platinum-based alkylating agent that induces DNA damage, has been shown to trigger cell death via oxidative stress and ferroptosis.^46,48^ Whether deregulation of ABCB7 leads to cisplatin resistance in group 3 tumors or if miR-1253 can sensitize cisplatin response through inhibition of ABCB7 remains unstudied. Here, we show that targeting ABCB7 can potentiate cisplatin cytotoxicity in G3MB by inducing iron overload, oxidative stress, and lipid peroxidation, which together trigger ferroptosis.

## Materials and Methods

### Patient Samples

Formalin-fixed paraffin embedded tissue blocks and frozen tissues of normal cerebellum (pediatric=12, adult=5) and pediatric MB specimens (WNT=1, SHH=9, grp 3=11, grp 4=16, unknown=7) were obtained from the Children’s Hospital and Medical Center, Omaha and the University of Nebraska Medical Center after Institutional Review Board approval. Informed consent was not required since the status of the study was exempted. For expression profiles of ABCB7 and GPX4, we cross-analyzed two primary MB datasets (Kanchan *et al.*, GSE148390 and Weishaupt *et al*., GSE124814).^49–51^ For Spearman correlation, we used GSE148390. For Kaplan-Meier Survival Analysis, we used the R2 database (Cavalli *et al.*, GSE85217).^9^

### Cell Lines and Cell Culture

D283 and D341 were purchased from ATCC (Manassas, Virginia); D425 and D556 were kind gifts from Darell Binger (Duke University Medical Center, Durham, NC); HDMB03 cells were a kind gift from Till Milde (Hopp Children’s Tumor Center, Heidelberg, Germany). Cell line genotyping was verified using short tandem repeat (STR) DNA profiling (UNMC). D283, D341, D425 and D556 cell line were maintained in DMEM supplemented with 10%-20% FBS and 100µg/ml penicillin/streptomycin. Normal human astrocytes (NHA) were purchased from Lonza Bioscience (Walkersville, MD) and grown in ABM basal medium supplemented with growth factors (Lonza Biosciences). All cell lines were maintained in 95% humidity, 37^D^C, 5% CO_2_.

### Transient Transfections

Cells at a density of 0.5 x 10^6^ were seeded in 6-well plates for 24 h and subsequently serum starved for 4 h prior to transfection. Cells were transfected with miR-1253 mimic (miRVanaTM miRNA mimic, ThermoFisher, 100 nM) or scramble negative control (100 nM) with Lipofectamine 2000 (Invitrogen) for 24 h.

### CRISPR/Cas9 Knockouts

Lentiviral particles were prepared by transfection of plasmid expressing Cas9 or sgRNA of ABCB7 (Addgene) co-transfected with pCMV-dR8.2 dvpr (Addgene) and pCMV-VSV-g (Addgene) lentiviral packaging plasmids into HEK293T cells using polyethyleneimine (PEI) transfection reagent. Virus-containing supernatant was collected and filtered 48 h after transfection. HDMB03 cells in 6-well plates were infected with Cas9 viral supernatant containing 4 μg/mL polybrene. Following blasticidin selection (10 μg/ml), the expression of Cas9 was confirmed by Western blotting. Stable HDMB03 cells expressing Cas9 expression were infected with the ABCB7 sgRNA viral supernatant containing 4 μg/ml polybrene. After 24 h of infection, cells were selected with 0.5 μg/ml puromycin. Single cell clones with Cas9 expression and ABCB7 knockout were amplified and used for subsequent experiments.

### Cell Viability

After transfection, HDMB03 cells (5 x 10^3^ cells/well) were re-seeded into 96-well plates and treated with defined concentrations of cisplatin (1-50 μM) for 24-72 h. Subsequently, MTT (5 mg/mL) or XTT (0.3 mg/mL) was added to each well and incubated for 2 or 6 h, respectively, at 37 °C. MTT absorbance was measured at 570 nm; XTT absorbance was measured at 440 nm. Data were analyzed using the SOFTMAX PRO software (Molecular Devices Corp., Sunnyvale, CA, USA).

### Colony Formation

After transfection, HDMB03 cells (1 x 10^3^ cells/well) were re-seeded in 6-well plates. Cells were treated with cisplatin (2 µM) and grown in complete medium for 9 days. Colonies were stained with 0.25% crystal violet (dissolved in 50% methanol) for 30 min. Crystal violet was dissolved in 10% acetic acid and absorbance read at 590 nm.

### Luciferase Assay

Luciferase assay was performed as previously described.^27^ Primers ABCB7 Forward: 5’-TAAGCCTGACATAACGAGGA-3’; ABCB7 Reverse: 5’-GCATCTCAGTATTAACTCTAGC -3’) were purchased from Eurofins. 3’UTR Wild and 3’UTR-Mutant were incorporated into XbaI restriction site of PGL3-control vector (Promega) expressing firefly luciferase. Dual luciferase assay was done in HDMB03 cells (3 x 10^5^cells/well) in 12-well plates. Luciferase activity was the measured using Dual-Luciferase Reporter Assay System (Promega) with a Luminometer (Biotek).

### Calcein AM Staining

Calcein-acetoxymethyl ester (Calcein AM) is a membrane permeable, non-fluorescent dye which emits green fluorescence once internalized into the live cells and cleaved by cytoplasmic esterase. Although calcein fluorescence is stable, it can be quenched by divalent metal ions such as iron and cobalt. To estimate cytosolic labile iron pool (LIP), HDMB03 and D425 cells were either transfected with scramble or miR-1253 or had ABCB7 knocked down. Cells were reseeded on to glass coverslips treated with or without the iron chelator deferoxamine (DFO) for 6 h. Cells were washed twice with 0.5 ml of PBS and incubated with 0.05 μM Calcein AM for 15 min at 37°C. Cells were analyzed under confocal microscope. Ex/Em = 488 nm/525 nm.

### Detection of Cytosolic and Mitochondrial Fe^2+^

Scramble vs. miR-1253-transfected cells or wild-type vs. ABCB7^KO^ cells were seeded onto glass coverslips and were treated with or without DFO (100 µM) for 6 h. For cytosolic LIP, 1 μmol/L FerroOrange (Dojindo, Japan) and 100 nM MitoTraker^TM^ Deep Red FM were added to each well; for mitochondrial LIP, 1 µM/L Mito-FerroGreen (Dojindo, Japan) and 100 nM MitoTraker Deep Red FM were added to each well. Cells were incubated in a 37° C incubator equilibrated with 95% air and 5% CO_2_ for 30 min. After incubation cells were washed and counter-stained with 4′,6-diamidino-2-phenylindole (DAPI). Cells were observed under a confocal fluorescence microscope, Ex/Em_cytosolicLIP_ = 561 nm/570-620 nm, Ex/Em_mitochondrialLIP_ = 488 nm/500-550 nm.

### Measurement of Intracellular Oxidative Stress

Intracellular ROS (H_2_O_2_) was measured using oxidation-sensitive fluorescent probe 2′,7′-Dichlorofluorescin diacetate (DCFDA) (Sigma Aldrich, USA); mitochondrial O_2_ ^•−^ was measured using MitoSOX™ Red (Thermofisher, USA). Scramble vs. miR-1253-transfected cells or wild-type vs. ABCB7^KO^ cells were treated with cisplatin or cisplatin and Ferrostatin-1 (Apexbio, USA) for 24 h. Cells were incubated with 10 µM DCFDA or 5 µM MitoSOX™ Red for 30 min. Oxidized DCFDA and MitoSOX™ Red were measured using at Ex/Em ∼485/528 nm and Ex/Em ∼510/580, respectively.^52^ Images were captured using EVOS FL Auto Imaging System (EVOS FL Auto, Life Technologies). Oxidized DCFDA and MitoSOXRed were measured using multimode plate reader at Ex/Em ∼485/528 nm and Ex/Em ∼510/580, respectively

### Measurement of Lipid Peroxidation

The Image-iT® LPO kit was used to measure lipid ROS through oxidation of the C-11-BODIPY® 581/591 sensor according to the manufacturer’s instructions. Briefly, scramble vs. miR-1253-transfected cells or wild-type vs. ABCB7^KO^ cells were treated with or without cisplatin or a combination of cisplatin and Ferrostatin-1 for 24 h. Cells were then stained with Image-iT® Lipid Peroxidation Sensor (10 μM) for 30 min and counter-stained with DAPI. Images were captured by confocal microscopy.

### Ferroptosis Assessment by Flow Cytometry

Scramble and miR-1253-transfected cells were treated with or without cisplatin or a combination of cisplatin and Ferrostatin-1 for 24 h. After completion of treatment, cells were incubated with 5 µM of MitoSOX™ Red for 30 min. Cells were washed and stained with Annexin-V/Cy™5(BD Biosciences). Cell populations were sorted and measured by flow cytometry.

### Glutathione Estimation

Glutathione estimation was performed according to the manufacturer’s instructions using GSH-Glo™ Glutathione Assay kit (Promega, USA). Briefly, scramble vs. miR-1253-transfected cells or wild-type vs. ABCB7^KO^ cells were seeded (5× 10^3^ cells/well) in the 96-well plates. Cells were treated with or without cisplatin or in combination with cisplatin and Ferrostatin-1 for 24 h. Plates were incubated in the dark for 30 min on a plate shaker with 100 µl of prepared 1x GSH-Glo™ Reagent. Then, 100 µl of reconstituted Luciferin Detection Reagent was added to each well and incubated in the dark for another 15 min. A standard curve was prepared using GSH standard solution to facilitate the conversion of luminescence to GSH concentration.

### Immunofluorescence Imaging

In cultured cells: scramble vs. miR-1253-transfected cells or wild-type vs. ABCB7^KO^ cells were seeded onto coverslips, rinsed, and fixed using 4% paraformaldehyde (PFA) in PBS for 10 min at room temperature. Cells were washed, incubated with 0.25% Triton X-100, and blocked with 3% BSA at room temperature. Cells were then incubated with primary antibody ABCB7 or COXIV overnight. Following day, cells were incubated with Alexafluor 488 and Alexaflour 547 conjugated antibodies for 1 h. Cells were washed and counter stained with DAPI. Images were captured at 63X using Carl Zeiss microscope (LSM 800 META).

In tissue: immunofluorescence was performed on surgically-resected, formalin-fixed, paraffin-embedded sections of group 3 medulloblastomas. Deparaffinized tissue sections were blocked with 5% BSA after heat induced epitope retrieval with citrate buffer (pH 6.0) and then incubated with primary antibodies, ABCB7 rabbit monoclonal antibody (1:200), GPX4 rabbit monoclonal antibody, or COXIV mouse monoclonal antibody (1:200). Following overnight incubation with primary antibody, sections were incubated for 1 h with Alexa-488 or Alexa-547-conjugated, mouse and rabbit secondary antibodies (1:200). Sections were counter-stained with DAPI. Micrographs were captured at 20X using Carl Zeiss microscope (LSM 800 META).

### PCR Primers

Total RNA extraction and quantitative PCR (qPCR) were performed according to manufacturer protocol. The forward (F) and reverse (R) primer sequences were as follows: ABCB7 (F): 5’-AAGATGTGAGCCTGGAAAGC-3’ (R): 5’-AGAGGACAGCATCCTGAGGT-3’; GPx4 (F): 5’-ACAAGAACGGCTGCGTGGTGAA-3’ (R): 5’-GCCACACACTTGTGGAGCTAGA-3’; β-Actin (F): 5’-CACCATTGGCAATGAGCGGTTC-3’(R): 5’-AGGTCTTTGCGGATGTCCACGT-3’.

### Statistical Analysis

Data are presented as mean ± SD. All experiments were conducted at least in duplicates with 3-6 replicates per experiment. Statistical analyses were performed using Prism 9.2 (GraphPad Software). Differences between groups were compared using Student’s t-test or one-way analysis of variance (ANOVA). Statistical significance was established at \**p* <0.05; \*\**p* <0.01; \*\*\**p* <0.001; \*\*\*\**p* <0.0001. Statistical analyses of high-throughput sequencing data were performed using R Statistical Software v4.1.1 (R Core Team), expression values were compared using a Mann-Whitney *U* test.

## Results

### Ferroptosis evasion characterizes G3MBs

Iron is vital to cellular processes, such as DNA synthesis, proliferation, cell cycle regulation, electron transport and catalysis, and oxygen transport and delivery.^53,54^ Rapidly dividing cancer cells can reprogram iron metabolism by upregulating essential iron regulatory and transport genes to support their invasive and rapid growth.^54^ Ferroptosis occurs when free ferrous iron oxidizes lipid membranes.^55^ Ferroptosis evasion, in turn, would result from mitigating iron imbalance and oxidative stress.

Using single sample gene set enrichment (ssGSEA) in a large MB dataset (GSE85217)^9^, we first observed that ferroptosis was the most enriched programmed cell death (PCD) pathway in MB tumors compared to normal **(Supplementar Figure 1A)**. We then compared the survival of patients in this cohort based on their ferroptosis scores, a measure of how active this pathway is. We observed that patients with lower ferroptosis scores (suggestive of lower activation) had lower survival **(Supplementary Figure 1B)**. Amongst the 4 MB subgroups, groups 3 and 4 have significantly lower survival than WNT and SHH tumors.^56^ And between these two subgroups, G3MBs had significantly lower ferroptosis scores than G4MB **(Supplementary Figure 1C)**. Our data suggest that lower ferroptosis activation (ferroptosis evasion) may be associated with more aggressive tumors.

When comparing G3MBs and normal cerebellum, we discovered enrichment of iron homeostasis (IREB2, STEAP3), iron transport (TFRC, ABCB6-8, SLC40A1, SLC25A37), glutathione metabolism (GCLC, GSS, TRIT1, SEPSECS), and ferroptosis regulatory genes that inhibitor ferroptosis. GPX4, the central regulator of ferroptosis^57^, was also enriched in our analysis **(Supplementary Figure 1D)**. We then queried the Molecular Signature Database (MSigDB, version 7.4) for upregulated pathways associated with the isolated genes. We discovered the enrichment of iron-sulfur (Fe-S) cluster binding and downregulation of ROS pathways **(Supplementary Figure 1E)**. Together, our data suggest that G3MBs, like many aggressive cancers, enrich iron homeostasis genes and redox pathways.^57,58^ This gene enrichment signature may enable G3MB tumors to counter iron imbalance and withstand oxidative stress and ferroptosis during their unchecked growth, enhancing their proliferative and aggressive phenotype.

### ABCB7, a novel target of miR-1253, is enriched in group 3 medulloblastomas

In our previous study, we identified miR-1253 as a novel tumor suppressor gene in MB, along with its multiple oncogenic targets.^27^ Upon restoration of miR-1253, using transient overexpression in HDMB03 cells, we observed potent downregulation of the ABC transporter superfamily by KEGG pathways analysis **(Supplementary Figures 2A)**. We wanted to then isolate specific targets of miR-1253 relevant to G3MB aggressiveness. We began by identifying ABC transporters enriched in G3MB **(Supplementary** Figure 2B**, column 1)**. From this list, we isolated transporters whose high expression conferred a poor prognosis **(Supplementary Figure 2B, column 2, red)**. From this cohort, we selected transporters whose expression was inhibited by miR-1253 overexpression in HDMB03 cells **(Supplementary** Figure 2B**, column 3, red)**. This comparative analysis revealed ABCB7.

ABCB7 resides on the inner mitochondrial membrane and plays a key role in Fe-S cluster biosynthesis and export.^59–61^ Notably, our transcriptomic analysis isolated Fe-S cluster binding as a top deregulated pathway and ABCB7 as a top enriched gene in G3MB tumors. In multiple MB cohorts, ABCB7 was shown to be upregulated **(Supplementary Figure 2C).** We then compared subgroup-specific ABCB7 expression in 3 MB cohorts and identified consistent enrichment of ABCB7 in G3MB (**Figure 1A**; **Supplementary Figures 2D and 2E)**. High ABCB7 expression in MB tumors was associated with dramatically poorer overall survival **(Supplementary Figure 2F)**. When the same survival analysis was run in a subgroup-specific manner, we learned that ABCB7 upregulation is associated with poor survival only in G3MB tumors (**Figure 1B**). These findings strengthened our premise to study ABCB7 enrichment as a possible mechanism for G3MB aggressiveness.

**Figure 1.**
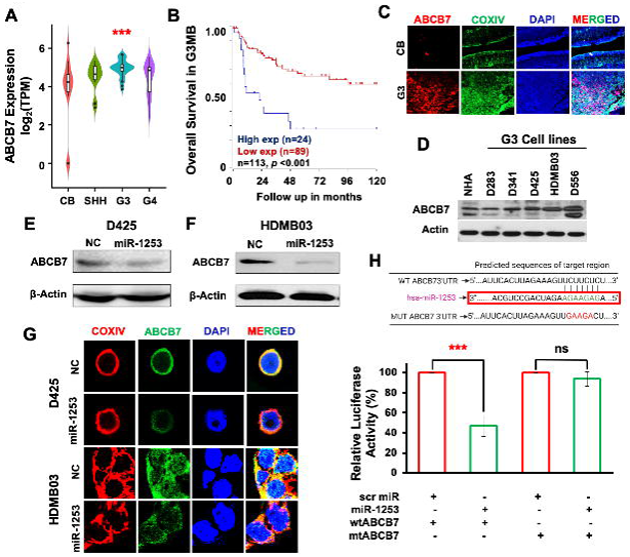
ABCB7 expression upregulation in G3MB portends a poor prognosis. **(A)** Subgroup-specific ABCB7 expression assessment by RNA sequencing (log_2_ transcripts per million) of a local medulloblastoma patient cohort (Kanchan *et al.*, GSE) showing specific upregulation in G3MB. *CB*, pediatric cerebellum (n=10); *SHH*, sonic hedgehog (n=6); *G3*, group 3 (n=7); *G4*, group 4 (n=12). **(B)** Poor prognostic profile demonstrable in high-expressing G3MB patients (Cavali *et al*. GSE85217). **(C)** Confocal microscopic images confirming high ABCB7 expression in G3MB tumors (n=6) compared to pediatric cerebellum (n=6) and colocalization to the mitochondria based on COXIV fluorescence. Images captured at 10X magnification. **(D)** Western blotting analysis showing high ABCB7 expression in classic MB cell lines (G3: D341, D425, HDMB03, D556; G3/4: D283) compared to normal human astrocytes (NHA). Downregulation of ABCB7 with miR-1253 overexpression in **(E)** D425 and **(F)** HDMB03 cells. **(G)** Confocal microscopic images in D425 and HDMB03 cells showing co-localization of ABCB7 to the mitochondria and downregulation with miR-1253 overexpression. Images captured at 63X magnification. **(H)** Dual-luciferase assay confirming direct binding of miR-1253 to ABCB7 in HDMB03 cells. Data presented as mean ± SD, n=3; *ns*, not significant, \*\*\**p* <0.001. *NC*, negative control; *Scr miR*, scramble micro RNA.

Now focusing on ABCB7, we first confirmed by confocal microscopy that ABCB7 co-localizes to the mitochondria, as evidenced by co-staining with COXIV, an inner mitochondrial membrane protein critical to electron transport (**Figure 1C**). Additionally, we confirmed robust expression of ABCB7 in a panel of aggressive group 3 MB cell lines compared to normal human astrocytes (**Figure 1D**).

To demonstrate that ABCB7 is inhibited by miR-1253, we first showed translational repression via Western blotting in 2 group 3 MB cell lines, D425 (**Figure 1E**) and HDMB03 (**Figure 1F**). In the same cell lines, we showed co-localization of ABCB7 with COXIV to the mitochondria and visualized the effect of miR-1253 on ABCB7 expression, as illustrated by a dramatically reduced green fluorescence by confocal microscopy (**Figure 1G**). Finally, we showed direct binding of miR-1253 to ABCB7 via a dual-luciferase reporter assay (**Figures 1H**). Together, these data confirmed that i) ABCB7 expression is enriched in group 3 tumors, ii) ABCB7 enrichment is a poor prognostic sign in group 3 patients, and iii) miR-1253 effectively inhibits ABCB7 expression.

### ABCB7 repression attenuates G3MB cancer cell aggressiveness

While the pathogenic role of ABCB7 has yet to be elucidated in MB, in glioma cells, ABCB7 inhibition triggered both apoptotic and non-apoptotic cell death through iron overload and ROS.^42^ Given the promiscuous nature of miRs, we adopted additional strategies to repress ABCB7 expression in G3MB cancer cells by using small-interfering RNA (si-ABCB7) or dox-inducible knockdown **(Supplementary Figure 2G)**. Using these strategies, we observed potent inhibition of primary medullosphere formation (**Figure 2A**) and reduced cancer cell viability by MTT assay (**Figure 2B**), all evidencing that ABCB7 repression attenuates aggressiveness of HDMB03 cells.

**Figure 2.**
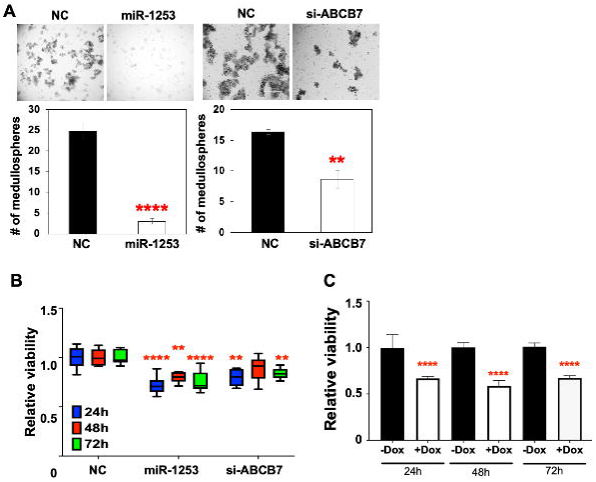
ABCB7 repression attenuates G3MB cancer cell aggressiveness. Attenuation of medullosphere formation **(A)** and reduction in viability (by MTT) of HDMB03 cells treated with miR-1253^OE^ (100 nM) and si-ABCB7 (50 nM). **(C)** Dox-inducible ABCB7^KD^ (1 ug/mL) demonstrating similarly reduced viability of HDMB03 cells. Data presented as mean ± SD, n=3; \*\**p* <0.01, \*\*\*\**p* <0.0001. *Dox*, doxycycline; *NC*, negative control.

### ABCB7 repression triggers iron overload

Given its primary role as a Fe-S cluster exporter, we studied the impact of repressing ABCB7 expression on iron balance in HDMB03 cells using ferric ammonium citrate (FAC) to mimic iron overload^62^ as our positive control. We observed an accumulation of free iron in the mitochondria (**MitoFerroGreen, Figure 3A and Supplementary Figure 3A**) and the cytoplasm (**FerroOrange, Figures 3B**, **3C, and Supplementary Figure 3B**) in HDMB03 cells with repressed ABCB7 expression. Iron chelation using deferoxamine (DFO) reverted the phenotype to wild-type. Calcein AM is a membrane-permeable dye that produces green fluorescence when internalized and can be rapidly quenched by divalent metals like iron. Repression of ABCB7 by miR-1253 led to quenching of Calcein AM in D425 **(Supplementary Figure 3C)** and HDMB03 cells **(Supplementary Figure 3D)**, further strengthening our observations of iron overload induced by ABCB7 inhibition.

**Figure 3.**
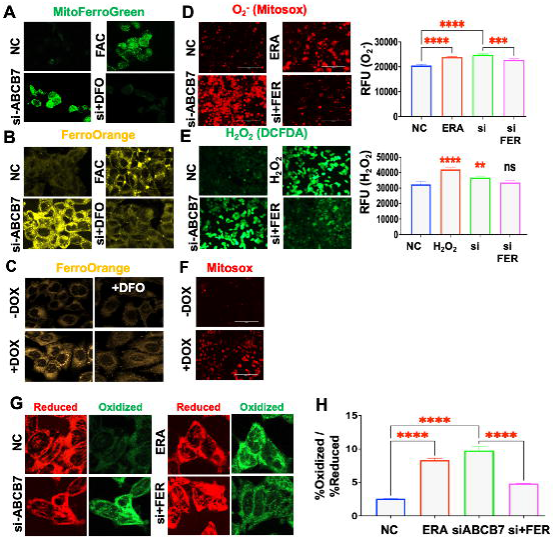
ABCB7 repression triggers iron imbalance, oxidative stress, and lipid peroxidation in G3MB cells. Iron overload demonstrable in the mitochondria **(A)** and cytoplasm **(B)** of HDMB03 transfected with si-ABCB7 (100 nM) as measured by MitoFerroGreen and FerroOrange, respectively. Deferoxamine (DFO) rescues this phenotype. **(C)** Cytoplasmic iron overload demonstrable with dox-inducible ABCB7^KD^ (1 ug/mL) in HDMB03 cells. Elevated mitochondrial **(D)** and total cellular **(E)** oxidative stress in HDMB03 cells transfected with si-ABCB7 (100 nM) by mitochondrial O ^•−^ (MitoSOX™ Red) and cellular H_2_O_2_ (DCFDA), respectively. Ferrostatin-1 (100 nM) rescues this phenotype. **(F)** Elevated mitochondrial oxidative stress demonstrable with dox-inducible ABCB7^KD^ (1 ug/mL) in HDMB03 cells. **(G)** Elevated lipid peroxidation in ABCB7-repressed HDMB03 cells as measured by BODIPY-C11 staining. **(H)** Lipid peroxidation quantified by FACS. Mean ±SD, n=3; *ns*, not significant, \*\**p*<0.01, \*\*\**p* <0.001 \*\*\*\**p*<0.0001. Fluorescent images for ROS obtained at 20X; quantification done by averaging 4 fields. Confocal images for lipid ROS obtained at 40X. Quantification of lipid ROS conducted via FACS, examining ratio of oxidized-to-reduced BODIPY-C11. *DFO*, deferoxamine; *ERA*, erastin; *FAC*, ferric ammonium citrate; *FER*, ferrostatin-1; *H_2_O_2_*, hydrogen peroxide; *O ^-^*, superoxide; s*i*, si-ABCB7.

### ABCB7 repression enhances cellular ROS and induces lipid peroxidation resulting in ferroptosis

Free ferrous iron (Fe^2+^) can convert to toxic ferric iron (Fe^3+^), via the Fenton reaction, which can disrupt the lipid bilayer to generate lethal lipid hydroperoxides, culminating in programmed cell death (PCD) by ferroptosis.^55,57^ Thus, after showing iron overload via ABCB7 repression, we subsequently quantified mitochondrial and total ROS in HDMB03 cells using hydrogen peroxide and erastin (ERA), a known inducer of ferroptosis^63^, as our positive controls. We observed elevated mitochondrial ROS (**Mitosox**, **Figure 3D and Supplementary Figure 4A**) and total ROS (**DCFDA**, **Figures 3E and Supplementary Figure 4B**) in ABCB7-repressed HDMB03 cells. We showed the same impact of ABCB7 repression by miR-1253 on oxidative stress in D425 cells **(Supplementary Figures 4C)**.

We next quantified lipid peroxidation through BODIPY-C11 staining, a membrane-targeted lipid ROS sensor, using an Image-iT^TM^ lipid peroxidation kit which provides a ratiometric indication of fatty acid oxidation that can be quantified by microscopy and flow cytometry (FACS). Repressing ABCB7 elicited lipid peroxidation in HDMB03 (**Figures 3G and 3H**) and D425 cells (**Supplementary Figure 4D)**.

Moreover, in the setting of mitochondrial iron overload and ROS, we studied the impact of repressing ABCB7 on mitochondrial membrane potential and oxidative phosphorylation capacity. Using tetramethylrhodamine methyl ester (TMRM), which fluoresces within mitochondria with intact membrane potentials, we showed a dramatic reduction in signal with si-ABCB7, suggesting that ABCB7 repression leads to mitochondrial dysfunction (**Figure 4A**).^64,65^ ABCB7 repression by miR-1253^OE^ also impaired the oxygen consumption rate (OCR) of HMDB03 cells, a direct measure of mitochondrial ATP production. It concurrently increased the extracellular acidification rate (ECAR), a measure of ATP production under anaerobic conditions by glycolysis (**Figure 4B**).

**Figure 4.**
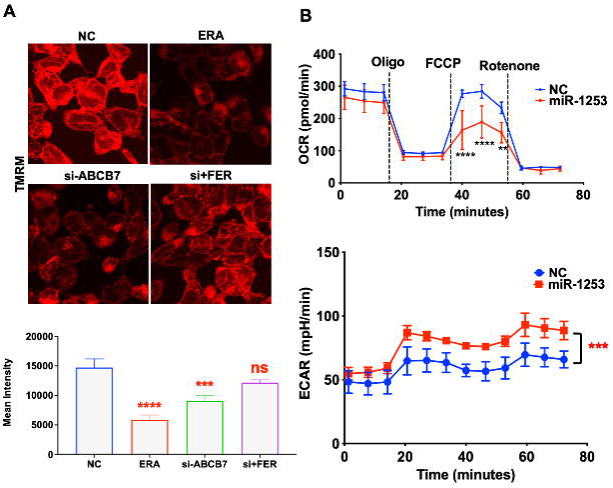
ABCB7 repression induces mitochondrial dysfunction. **(A)** Disruption of mitochondrial membrane potential as measured by TMRM fluorescence in ABCB7-repressed (si-ABCB7, 100 nM) HDMB03 cells, rescued by ferrostatin-1 (100nM). **(B)** Seahorse assay further showing disruption of oxidative phosphorylation in ABCB7-repressed (miR-1253 100 nM and si-ABCB7 100 nM) HDMB03 cells. Confocal images obtained at 63X magnification. Quantification done by taking average of 4 fields. Mean ±SD, n=3; *ns*, not significant, \*\*\**p*<0.001, \*\*\*\**p*<0.0001 compared to NC. *ERA*, erastin; *FER*, ferrostatin-1; *NC*, negative control; *si-*, si-ABCB7.

Ferroptosis is defined as an iron-dependent form of regulated cell death resulting from lipid peroxidation.^43^ Ferrostain-1 (FER) scavenges ROS and lipid hyperoxides and prevents ferroptosis.^66^ Given our observations of iron accumulation, ROS generation, and lipid peroxidation, we wanted to assess whether ABCB7 repression can, in fact, trigger ferroptosis. D425 and HDMB03 cells transfected with miR-1253 were stained with MitoSOX™ Red (for O_2_ ^•−^) and Annexin-V/Cy™5 (marker of apoptotic cell death). Flow cytometry analysis revealed a significant (∼3-fold) rise in dual-stained cells (Q2), indicating oxidative stress-mediated cell death **(Supplementary Figures 4E and 4F)**. Moreover, ferrostatin-1 rescued ABCB7-repressed cells from both oxidative stress (**Figures 3D**, **3E, and Supplementary Figures 4A and 4B)** and lipid peroxidation (**Figure 3G and 3H**). These are the <u>first data</u> showing that ABCB7 repression can trigger events that culminate in ferroptosis.

### ABCB7 repression inhibits G3MB tumor growth

We next examined the impact of ABCB7 repression on tumor burden in a validated G3MB mouse model^28^ using dox-inducible ABCB7 knockdown. We generated tumors in NOD *scid* gamma (NSG) mice by orthotopically injecting wildtype or ABCB7^KD^ HDMB03 cells (1x10^5^) into the cerebellum. We observed smaller tumors sizes in dox-induced ABCB7^KD^ mice at day 28 and day 30 (**Figures 5A and 5B**). This was concurrent with significantly lower immunohistochemical staining for ABCB7 and GPX4, as well as higher staining for 4HNE, a marker of lipid peroxidation (**Figure 5C**). These data reveal that ABCB7^KD^ attenuates G3MB tumor burden.

**Figure 5.**
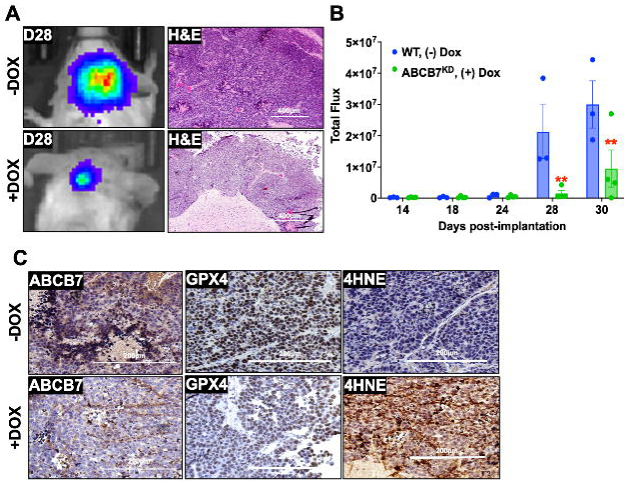
ABCB7 repression reduces G3MB tumor burden. NSG mice were orthotopically injected with luciferase-tagged wild-type vs. dox-inducible ABCB7^KD^ HDMB03 cells. Dox induction was started at day 10 post-implantation. **(A and B)** Reduction in tumor burden noted in dox-induced ABCB7^KD^ group at day 28 and day 30 post-implantation compared to control. **(C)** Attenuated expression of ABCB7 and GPX4 and enhanced expression of 4HNE, a stable end-product of ferroptosis, in dox-induced ABCB7^KD^ group by immunohistochemical staining. Mean ±SD, n=3-5 mice per arm; \*\**p*<0.01 compared to -Dox mice. *4-HNE*, 4-hydroxynonenol; *Dox*, doxycycline; *GPX4*, glutathione peroxidase 4.

### ABCB7 repression inhibits GPX4, the primary regulator of ferroptosis

Cancer cells have the inherent ability to activate redox buffering systems to thrive in a highly oxidative environment.^67^ Glutathione (GSH) is a key player in this response and a critical cofactor for glutathione peroxidase 4 (GPX4). GPX4 uses GSH to reduce toxic lipid hydroperoxides to non-toxic lipid alcohols and is thus a central regulator of ferroptosis, found in the nucleus, cytoplasm, and mitochondria **(Supplementary Figure 5A)**.^57^ In G3MB, ROS pathways are downregulated and concurrent with elevated expression of genes involved in glutathione metabolism, including GSS, GCLC, GPX4 **(Supplementary Figures 1D and 1E)**. GPX4 overexpression elevates aggressiveness in multiple cancers (breast, liver, GI, pancreatic).^68^ In MB, we noted significant enrichment of GPX4 in all MB tumor subtypes, with highest expression in G3MB **(Supplementary Figures 5B and 5C)**. By RNA Sequencing analysis, critical mediators of GSH biosynthesis (GSS, GCLC) and GPX function (SEPSECS, TIRT1) were also upregulated in G3MB **(Supplementary Figure 5D)**. Immunofluorescence studies revealed high expression of GPX4 in G3MB and co-localization of GPX4 to the mitochondria **(Supplementary Figure 5E)**. In classic G3MB cell lines, GPX4 expression was significantly elevated compared to normal human astrocytes **(Supplementary Figure 5F)**.

Given the pivotal role of GPX4 in ferroptosis mitigation, we studied the impact of ABCB7 repression on GPX4, expecting an upregulation to counter the oxidative stress induced by iron overload. Instead, we observed the opposite. While miR-1253^OE^ reduced GPX4 transcript and protein levels in HDMB03 (**Figure 6A**) and D425 (**Figure 6B**), ABCB7^KO^ (**Figure 6C**) and ABCB7^KD^ (**Figure 6D**) abrogated expression. In support, we noted a positive Spearman correlation (R=0.72, *p*<0.001) between GPX4 and ABCB7 (**Figure 6E**). We further explored this relationship using fractionation studies. While miR-1253^OE^ reduced GPX4 levels in the cytoplasm and mitochondria (**Figure 6F**), ABCB7^KO^ (**Figure 6G**) and si-ABCB7 (**Figure 6H**) again showed near abrogation of GPX4 expression from the cell. GSH, a main substrate of GPX4, was concurrently reduced in miR-1253-overexpressing D425 **(Supplementary Figure 6A**) and HDMB03 cells **(Supplementary Figure 6B**) as well as in si-ABCB7-treated HDMB03 cells **(Supplementary Figure 6C**). This is the <u>first time</u> a link has been proposed between ABCB7 and GPX4, providing us preliminary insight into the impact of the miR-1253/ABCB7 axis on GPX4 and the cellular response to ferroptosis.

**Figure 6.**
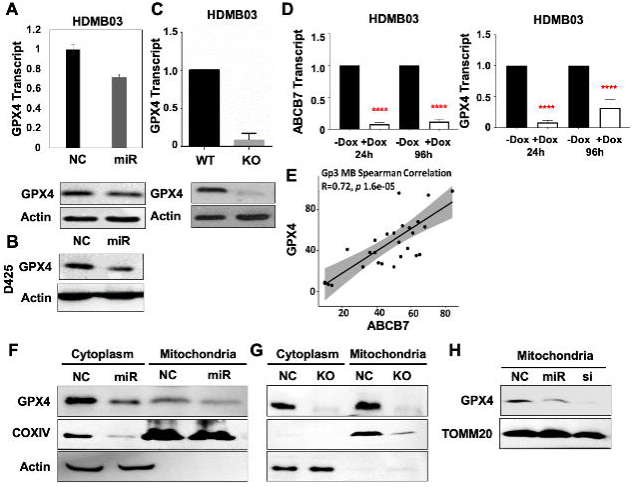
ABCB7 repression inhibits GPX4 expression, the central regulator of ferroptosis. Reduced GPX4 expression by miR-1253^OE^ in HDMB03 **(A)** and D425 **(B)**, by ABCB7^KO^ **(C)**, and ABCB7^KD^ **(D)**. **(E)** Spearman correlation showing positive correlation between ABCB7 and GPX4 expression. (Kanchan *et al.*, GSE148390). Fractionation studies revealing that ABCB7 repression by miR-1253^OE^ **(F)**, ABCB7^KO^ **(G)**, and si-ABCB7 **(H)** abrogates GPX4 expression from the cytoplasm and mitochondria. Mean ±SD, n=3; \*\*\*\**p*<0.0001. *KO*, ABCB7^KO^; *miR*, miR-1253 overexpression; *NC*, normal control; *si*, si-ABCB7; *WT*, wildtype HDMB03 cells.

### ABCB7 repression potentiates cisplatin cytotoxicity in G3MB cells

Cisplatin, a platinum-containing chemotherapeutic agent, induces DNA damage via various mechanisms, including (i) crosslinking DNA purine bases, (ii) inducing oxidative stress, and (iii) interfering with DNA repair machinery, that can eventually lead to cancer cell death.^69^ Recent data purport that it can also inhibit GSH and GPX4.^46,48,70,71^ In HDMB03 cells, we observed cisplatin to inhibit both ABCB7 and GPX4 expression (**Figure 7A**). In fact, we observed that cisplatin depleted GSH in HDMB03 cells with the addition of ferrostatin having some recovery effects. However, cisplatin added to cells with ABCB7 inhibition had the strongest effect in depleting GSH; ferrotstatin-1 catalyzed the recovery of reduced glutathione levels in these cells **(Supplementary** Figures 7D and 7E).

**Figure 7.**
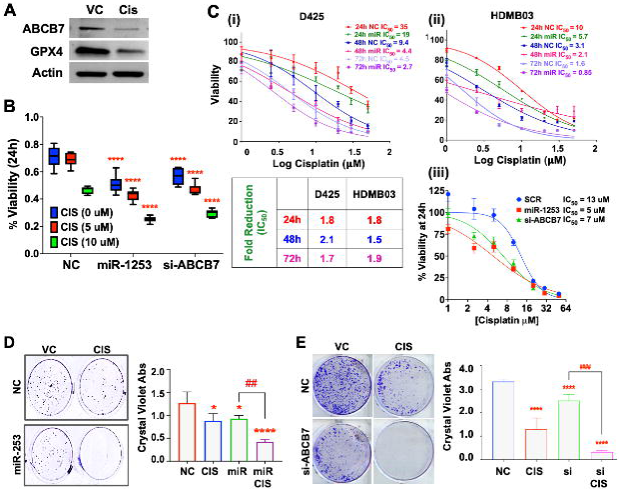
ABCB7 repression potentiates cisplatin toxicity in G3MB cells. **(A)** Cisplatin (2.5 μM) induced repression of ABCB7 and GPX4 expression in HDMB03 cells. **(B)** Potentiation of cisplatin toxicity in ABCB7-repressed HDMB03 cells by MTT. **(C)** A 2-fold reduction in IC_50_ of cisplatin in D425 **(i)** and HDMB03 **(ii)** cells transfected with scramble vs. miR-1253 for 24-72h and in HDMB03 cells transfected with si-ABCB7 (50 uM) for 24h **(iii)**. Tabulated results presented as fold-change in cisplatin IC_50_ between scramble and miR-1253 in D425 and HDMB03 cells at different time points. **(D and E)** Colonogenic assay demonstrating inhibitory effect of cisplatin vs. ABCB7 repression vs. combination on colony formation in HDMB03 cells. Mean ±SD, n=3; \**p*<0.05, \*\*\*\**p*<0.0001 compared to NC; ^##^*p*<0.01 compared to miR; ^####^*p*<0.001 compared to si. *CIS*, cisplatin; *FER*, ferrostatin-1; *miR*, miR-1253; *NC*, negative control; *si*, si-ABCB7.

In HDMB03 cells with repressed ABCB7 expression, cisplatin elicited higher cytotoxicity compared to controls (**Figure 7B**). Moreover, in D425 (**Figure 7Ci**) and HDMB03 (**Figure 7Cii**) cells, ABCB7 repression via miR-1253^OE^ resulted in a ∼2-fold reduction in the IC_50_ of cisplatin, while si-ABCB7 showed the same effect in HDMB03 cells (**Figure 7Ciii**). Colony formation of HDMB03 cells was nearly abrogated in ABCB7-repressed HDMB03 cells treated with cisplatin. (**Figure 7D**).

We next investigated whether cisplatin treatment enhanced total cellular and mitochondrial ROS in G3MB cells. We studied ABCB7 repression by miR-1253^OE^-transfected cells subjected to 10 µM cisplatin in D425 and 2 µM cisplatin in HDMB03 cells at 24 h; we concurrently examined cisplatin treatment (2 µM) of ABCB7^KO^ HDMB03 cells. As prior, ABCB7 repression by these methods induced significantly higher levels of total and mitochondrial ROS compared to control. Moreover, in all cases, ABCB7 repression potentiated cisplatin-mediated ROS (**Figures 8A-C**). Concurrent treatment with ferrostatin-1 returned ROS levels to that of untreated miR-1253^OE^ or si-ABCB7 (**Figure 8D and 8E**). In ABCB7KO HDMB03 cells, these effects were similarly recapitulated **(Supplementary Figures 8A and 8B).** These data provide evidence for cisplatin-induced cytotoxicity in G3MB cells by ferroptosis.

**Figure 8.**
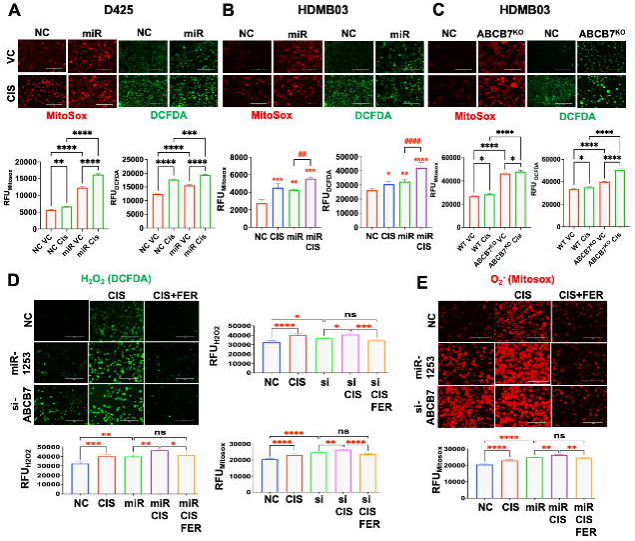
ABCB7 repression potentiates oxidative stress in cisplatin-treated G3MB cells. MiR-1253-transfected D425 **(A)** and HDMB03 **(B)**, and ABCB7^KO^ HDMB03 cells **(C)** treated with cisplatin (D425 24-h IC_25_ ∼10 µM; HDMB03 24-h IC_25_ ∼2 µM) demonstrating potentiated mitochondrial ROS (MitoSOX™ Red, O ^•−^) and total ROS (DCFDA, H O). **(D** and **E)** Cisplatin potentiating ROS in HDMB03 cells with repressed ABCB7 expression; ferrostatin-1 rescues this effect. Representative images and graphs showing the potentiating effect of combining miR-1253 with cisplatin. Images captured at 20X magnification. Scale bar 400 µm. Mean ± SD, n=3; *ns*, not significant, \**p*<0.05, \*\**p*<0.01, \*\*\**p*<0.001, \*\*\*\**p*<0.0001. *CIS*, cisplatin; *FER*, ferrostatin-1; *miR*, miR-1253; *NC*, negative control; *si*, si-ABCB7.

### ABCB7 repression potentiates cisplatin toxicity through ferroptosis in G3MB cells

We next examined the impact of cisplatin on lipid peroxidation to determine if cisplatin potentiated ferroptosis in ABCB7 repressed G3MB cells. First, cisplatin elicited a high degree of lipid peroxidation compared to erastin (**Figure 9A**). When combined with ABCB7 repression, cisplatin treatment potentiated lipid peroxidation. Treatment with ferrostatin-1 significantly reduced lipid peroxidation in cisplatin-treated ABCB7-repressed HDMB03 cells (**Figure 9B and 9C**).

**Figure 9.**
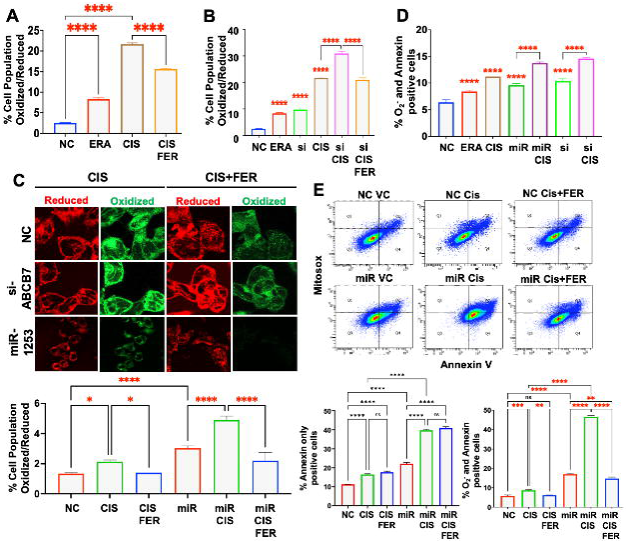
ABCB7 repression potentiates cisplatin cytotoxicity in G3MB cancer cells by triggering ferroptosis. Lipid peroxidation was visualized by BODIPY-C11 staining and quantified by FACs using the ratio of oxidized-to-reduced lipids in HDMB03 cells. **(A)** Cisplatin (5 uM) induced lipid peroxidation compared to erastin, a known inducer of ferroptosis, with partial rescue by ferrostatin-1. **(B)** Potentiation of lipid peroxidation in ABCB7-repressed cells treated with cisplatin (5 uM), with partial rescue by ferrostatin-1. **(C)** Visualization of lipid peroxidation as measured by BODIPY-C11 staining by confocal microscopy. Images obtained at 63X. **(D and E)** Analysis of cell death by flow cytometry showing significantly higher O_2_^•−^ mediated cell death (representing ferroptosis) in ABCB7-repressed HDMB03 cells. This is demonstrable by quantifying cells staining for both Annexin V-Cy5 (apoptosis) and for O ^•−^ (Mitosox) (Q2). ABCB7 repression by si-**(D)** and miR-1253 **(E)** potentiated cisplatin-induced ferroptosis, rescue by ferrostatin-1. Mean ± SD, n=3; \**p* <0.05, \*\**p* <0.01, \*\*\**p* <0.001, \*\*\*\**p* <0.0001. *CIS*, cisplatin; *ERA*, erastin; *FER*, ferrostatin-1; *miR*, miR-1253; *NC*, negative control; *si*, si-ABCB7.

Finally, examining O_2_-mediated cell death (as a proxy for ferroptosis) at 24 h, cisplatin showed a small rise in ROS-mediated cell death and apoptosis, while miR-1253 led to a larger rise in both. Treating miR-1253-transfected HDMB03 cells with cisplatin elicited a dramatic increase in both ROS-mediated and apoptotic cell death; ferrostatin-1 treatment rescued the former, strongly implicating the important contribution of ferroptosis to the mechanism of miR-1253-induced cell death (**Figure 9E**). Together these data substantiate the cardinal role of ABCB7 repression in inducing ferroptosis and its potentiating effects on cisplatin cytotoxicity through ferroptosis.

### Artesunate synergizes with cisplatin to kill G3MB cancer cells

Artesunate is an anti-malarial drug that kills *Plasmodium falciparum* by interacting with free iron to generate ROS.^72^ Its anti-neoplastic action has been demonstrated in head and neck cancers, where it reduced GSH levels, increased lipid ROS, and triggered ferroptosis.^73^ Moreover, tumor susceptibility to artesunate was enhanced by iron overload, especially in high TFRC-expressing tumors.^74^ In the same study, artesunate was shown to inhibit ABCB7 in human leukemia and breast cancer cells.^74^ These features make artesunate an ideal drug to test in G3MBs, which are also high TFRC-expressing tumors.

We studied the cytotoxicity of artesunate in HDMB03 cells. Our dose-response curves revealed a declining IC_50_ over 72h, suggestive of a cumulative effect over time (**Figure 10A**). Of note, these IC_50_ values are far lower than what has been determined for artesunate in glioma cell lines.^75^ Then, in orthotopic G3MB tumors developed in NSG mice, artesunate monotherapy was shown to effectively inhibit tumor growth at 27 days post-implantation and even marginally enhance survival (**Figure 10B**).

**Figure 10.**
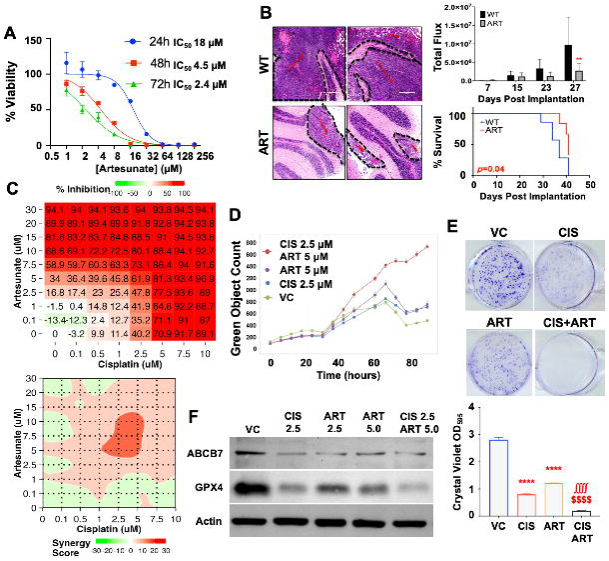
Artesunate synergizes with cisplatin to kill G3MB cancer cells. **(A)** Dose-response curve for artesunate in HDMB03 cells. **(B)** Reduction in G3MB tumor burden and prolonged survival with artesunate (100 mg/kg i.p. daily) in an orthotopic G3MB tumor model. n=3-5 mice per group. **(C)** Drug synergy experiment between cisplatin and artesunate showing synergy between both drugs in HDMB03 cells. Potentiated cell death **(D)** and abrogated colony formation **(E)** in with combined artesunate + cisplatin. **(F)** Potent inhibition of GPX4 by artesunate and cisplatin treatments. Mean ±SD, nb=3; \*\*\*\**p*<0.0001 compared to VC, ^∬∬^*p*<0.0001 compared to CIS, ^$$$$^*p*<0.0001 compared to ART. *ART*, artesunate; *CIS*, cisplatin; *FER*, ferrostatin-1; *GPX4*, glutathione peroxidase 4; *VC*, vehicle control.

Given our prior findings of ABCB7 repression potentiating cisplatin cytotoxicity, we tested whether artesunate can bolster cisplatin cytotoxicity by potentiating iron overload and ferroptosis. By performing a drug synergy experiment, we discovered that artesunate synergizes with cisplatin (highest synergy score 15.2) (**Figure 10C**). Combination therapy was superior to monotherapy in inducing cell death in HDMB03 cells (**Figure 10D**) and abrogated colony formation (**Figure 10E**). Finally, we observed, similar to cisplatin, that artesunate (5 μM) inhibited both ABCB7 and GPX4 in G3MB cells. However, combination therapy with cisplatin (2.5 μM) compounded GPX4 inhibition (**Figure 10F**). These data suggest that artesunate potentiates cisplatin cytotoxicity in G3MB cells and may serve as an innovative treatment strategy for G3MB tumors.

### Artesunate synergizes with cisplatin to mitigate G3MB tumor burden

We then tested the impact of combination therapy in our orthotopic G3MB mouse model generated in NSG mice from HDMB03 cells. We divided our groups into negative control (PBS), cisplatin i.p. (2.5 mg/kg), artesunate i.p. (100 mg/kg)^76^, and artesunate+cisplatin. Mice were randomized and treatment was started at day 10 post implantation (**Figure 11A**). At termination of the experiment, at day 22 when all wildtype mice had died, mice in the combination arm had significantly smaller tumors and higher survival compared to other groups (**Figure 11B**).

**Figure 11.**
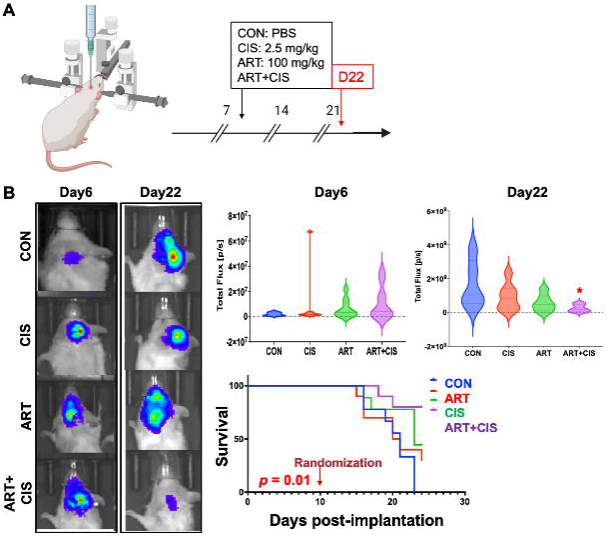
Artesunate synergizes with cisplatin to reduce G3MB tumor burden. **(A)** NSG mice were orthotopically injected with luciferase-tagged wild-type HDMB03 cells. Mice were randomized into four treatment arms (CON: negative control; CIS: cisplatin only at 2.5 mg/kg i.p.; ART: artesunate only at 100 mg/kg i.p.; ART+CIS: combination). Treatment was started at day 10 post-implantation. Experiment was terminated at day 22 when all wildtype mice died. **(B)** Mice in the combination arm had significantly smaller tumors and higher survival. Mean ± SD, n=3-5 mice per arm; \**p*<0.05 compared to PBS-treated negative controls. *ART*, artesunate; *ART+CIS*, combination arm; *CIS*, cisplatin; *CON*, control.

## Discussion

Amongst the most devastating diagnoses in a pediatric patient is a tumor of the central nervous system, with medulloblastoma being the most common malignant tumor.^3,4,77,78^ Poor survival (5-year OS <50%) in G3MB patients is attributable to a combination of young age at diagnosis (peak age 3-5), metastases at diagnosis (up to 50%), and c-Myc amplification.^9,28,79–82^ Current treatment regimens have yet to impact the dismal prognosis with stagnant survival rates seen over the last decade.^83^ An inability to tolerate mainstay of therapy especially for young patients fuels high relapse rates (∼30%) which are universally fatal. For example, in children under the age of 4 years relapse was noted in ∼60% of patients who did not receive upfront craniospinal irradiation. Moreover, relapse rates were highest in group 3 MB tumors and those with i17q and c-Myc amplification.^84^ In those that do manage to survive, irreversible damage to the hypothalamic-pituitary axis is sustained from cytotoxic treatment regimens resulting in short stature, cognitive impairments, and emotional lability.^85,86^ Thus, the need for novel therapeutic strategies that mitigate drug-related cytotoxicity yet accomplish widespread tumor abstraction for this subgroup is dire.

Elevated recurrence rates have often been conjectured to be linked to mechanisms for drug resistance in group 3 MB, which may include high expression of multi-drug resistance (MDR) genes belonging to the ABC transporter family.^37,38,40,87,88^ Cytotoxic drugs, in combination with tumor suppressive miRNAs, have the potential for a more complex and profound effect on tumorigenesis and may possess the ability to address drug resistance patterns, especially if they can target MDR genes.^89^ Here, we demonstrated a strong negative enrichment of the ABC transporter family with miR-1253 expression restoration in group 3 MB cells. Of the multiple transporters identified via our bioinformatics approach, we isolated ABCB7, whose deregulated expression profile in group 3 tumors was strongly associated with poor prognosis, as a target of miR-1253.

ABCB7 is an iron transporter residing on the inner mitochondrial membrane and involved in Fe-S cluster biogenesis.^59,90^ ABCB7 deficiency can lead to decreased expression of electron transport chain (ETC) complex proteins I, II, IV and V, which can trigger impaired oxidative phosphorylation and mitochondrial membrane integrity resulting in oxidative stress.^41,91–93^ In glioma cells, inhibition of ABCB7 resulted in disruption of iron transport and ROS generation triggering apoptotic and non-apoptotic cell death.^42^ Iron accumulation can trigger oxidative stress and vice versa^94,95^, resulting in lethal lipid peroxidation activating ferroptosis, which is distinct from apoptosis, necrosis, autophagy, and pyroptosis.^96^

With a strong premise for exploring iron imbalance through ABCB7 repression, our subsequent studies substantiated a role for this strategy in activating ferroptosis in G3MB. First, miR-1253 induced intracellular iron accumulation in D425 and HDMB03 cells, effectively abrogated by pre-treatment with an iron chelator, DFO. ABCB7^KO^ resulted in a similar phenotype. This resulted in elevated mitochondrial ROS (O_2_ ^•−^) and cytosolic ROS (H O). We also recorded a high lipid oxidation profile in these cell lines concurrent with miR-1253 expression. Finally, using AnnexinV and MitoSOX™ Red staining, we revealed a significantly elevated dual staining population of cells in miR-1253-expressing cells. Taken together, these results are highly indictive of ferroptosis induction by miR-1253 in group 3 MB cells.

Ferroptosis can also be induced by disruption of glutathione synthesis or inhibition of glutathione peroxidase 4.^97^ Cancer cells have the capacity to activate redox buffering systems to survive in a highly oxidative environment resulting from deregulated cellular functions.^67^ Glutathione is a key player in this response and a critical cofactor for glutathione peroxidase 4 (GPX4). GPX4, a central regulator of ferroptosis upregulated in various cancers, uses glutathione to reduce ROS and lipid hydroperoxide levels thus facilitating tumor cell survival in an environment with high oxidative stress.^98^ Additionally, GSH binds Fe^2+^ to prevent iron oxidation and is thus a critical component controlling the labile iron pool.^99^ Of note, ABCB7 harbors a GSH binding pocket, and GSH is a required substrate for cytosolic and nuclear Fe-S protein biogenesis and iron homeostasis.^61,100^ In group 3 tumors, we not only showed deregulated GPX4 expression but also a strong positive correlation with ABCB7. Resultantly, miR-1253 expression or ABCB7^KO^ strongly inhibited GPX4 expression and reduced glutathione levels. These data demonstrate the contribution of disrupting GPX4 and glutathione metabolism in miR-1253-mediated oxidative stress mechanisms resulting in ferroptosis.

MiRNA mimics have been shown to possess the capability of restoring the sensitivity of cancer cells for chemotherapeutic agents and to thus subsequently enhance their effectiveness. For example, miR-429, miR-383, miR-101-3p, miR-195, miR-634, and miR-1294 elicited a 2-5 fold reduction in the EC_50_ and IC_50_ values in combination with gemcitabine, temozolomide, and paclitaxel.^89,101–106^ Of the standard chemotherapies for medulloblastoma tumors, only cisplatin has been shown to trigger both apoptosis and ferroptosis, via oxidative stress, GSH depletion, and GPX4 inactivation.^46,48,70,71^ Moreover, a microarray-based study of the IC_50_ of cisplatin in 60 NCI cell lines identified multiple ABC transporters, including 3 iron transporters, i.e. ABCB6, ABCB7 and ABCB10, in conferring cisplatin drug resistance.^107^

Given the ferroptotic mechanism we elucidated for miR-1253, we studied whether miR-1253 expression can potentiate cisplatin cytotoxicity in group 3 MB cells. We report a 2-fold reduction in the IC_50_ value of cisplatin with miR-1253 induction in D425 and HDMB03 cells. Combination treatment had dramatic effects on the clonal potential of HDMB03 cells. We then demonstrated the highest induction of mitochondrial and cytosolic ROS, lipid peroxidation, and GSH depletion in miR-1253-expressing HDMB03 and ABCB7^KO^ HDMB03 cells treated with cisplatin. Consequently, combination therapy resulted in the highest degree of ferroptosis in these cell lines. These effects were reversed by ferrostatin-1, a potent inhibitor of ferroptosis, lending further validity to the central role of ferroptosis in the mechanism of miR-1253-mediated effects and cisplatin potentiation.

Overall, our study has identified novel tumor suppressive properties for miR-1253. First, miR-1253 directly inhibits ABCB7 expression, thus inducing labile iron pool within MB cancer cells and stimulating ROS production. MiR-1253 can concurrently downregulate the expression of GPX4 and deplete GSH, further exacerbating ROS. Together, the generation of lipid hyperoxides progresses unabated, leading to cancer cell death via ferroptosis. We also leveraged these properties by showing potentiation of cisplatin cytotoxicity and thus enhanced therapeutic efficacy in group 3 MB cells. Together, our findings provide proof-of-concept for further exploration of tumor suppressive microRNAs as therapeutic adjuncts to standard chemotherapy. Such a strategy may mitigate the current limitations to treatment regimens in our youngest high risk patients.

## Author contributions

RKK and SM conceptualized and designed the study. RKK, JBK, NP, DD, and PG performed all experiments. PK developed ABCB7 knockout clone for the study. DD, RCV, and IT provided bioinformatics support for the study. SM coordinated and supervised data analysis. RKK wrote the first draft of the manuscript. NP, PK, DD, PG, RCV, JAS, RV, MWN SKB and SM provided scientific feedback, critical reviews, and helped in revision of the manuscript. All authors have read and approved the final manuscript.

## Abbreviations

ABC: ATP binding cassette
ANOVA: analysis of variance
BLC-2: B-cell lymphoma 2
BRCA1: breast cancer gene 1
CB: cerebellum
CD276: cluster of differentiation 276 (B7-H3)
CDK6: cyclin-dependent kinase 6
c-Myc: c-myelocytomatosis oncogene
DDR: DNA damage repair
DCFDA: 2′,7′-Dichlorofluorescin diacetate
DFO: deferoxamine
FACs: fluorescence-activated cell sorting
FISH: fluorescence *in situ* hybridization
GPX4: glutathione peroxidase 4
GSH: glutathione
GSS: glutathione synthetase
IC_50_: 50% inhibitory concentration
IHC: immunohistochemistry
i(17q): isochromosome 17q
KEGG: Kyoto encyclopedia of genes and genomes
KO: knock-out
LIP: labile iron pool
LPO: lipid peroxidation
MB: medulloblastoma
MDR: multiple drug resistance
miR: microRNA
miR-1253: microRNA 1253
MnTBAP: Mn(III)tetrakis(4-benzoic acid)porphyrin
mtROS: mitochondrial reactive oxygen species
MTT: 3-(4,5-dimethylthiazol-2-yl)-2,5-diphenyltetrazolium bromide
NC: negative control
NHA: normal human astrocytes
Non-SHH/WNT: non-Sonic Hedgehog/non-Wingless
PARP: poly ADP ribose polymerase
PCR: polymerase chain reaction
Ped: pediatric
PNET: primitive neuro-ectodermal tumor
ROS: reactive oxygen species
RSL3: RAS-selective lethal
SHH: Sonic Hedgehog
WNT: Wingless
XIAP: X-linked inhibitor of apoptosis
XTT: sodium 3′-[1-[(phenylamino)-carbony]-3,4-tetrazolium]-bis(4-methoxy-6-nitro)benzene-sulfonic acid hydrate

---

**Supplementary Figure 1. Iron homeostasis and glutathione metabolism are deregulated in G3MB. (A)** Genes associated with programmed cell death highlighting ferroptosis as the top deregulated pathway in MB patients (GSE82517). **(B)** Survival disadvantage for MB patients with lower ferroptosis scores. **(C)** G3MB tumors have lower ferroptosis scores than G4MBs. **(D)** Key genes regulating iron homeostasis, glutathione metabolism, and ferroptosis that are enriched in G3MB (GSE148389, *p*<0.05 for all presented gene comparisons between normal and G3MB). **(E)** Enrichment of iron-sulfur cluster binding and downregulation of ROS pathways in G3MB tumors.

**Supplementary Figure 2. ABCB7 is enriched in MB and portends a poor prognosis. (A)** Enrichment plot by KEGG pathways analysis demonstrating a strong negative enrichment score for the ABC transporter family with miR-1253 overexpression in HDMB03 cells. **(B)** Deregulated ABC transporters in G3MB (column 1), effect of deregulated expression on survival (column 2), and transporters significantly downregulated by miR-1253 expression (column 3). This analysis isolated ABCB7 as the best putative target for miR-1253. **(C)** Expression profile for ABCB7 in multiple MB cohorts. *CB*, normal cerebellum, Roth *et al.* 2008 (n=9, GSE3526); *MB 1*, Gilbertson *et al.* 2012 (n=76, GSE37418); *MB 2*, Pfister *et al.* 2017 (n=223); *MB 3*, Delattre *et al.* 2012 (n=57); *MB 4*, Kool *et al.* 2009 (n=62, GSE10327). Subgroup-specific ABCB7 expression in two separate medulloblastoma patient cohorts, i.e. **(D)** Weishaupt *et al.*, GSE124814; *CB*, normal cerebellum (n=291); *WNT*, wingless (n=118); *SHH*, sonic hedgehog (n=405); *G3*, group 3 (n=233); *G4*, group 4 (n=530); and **(E)** Luo *et al.*, GSE164677; *CB*, normal cerebellum (n=4); *WNT*, wingless (n=6); *SHH*, sonic hedgehog (n=20); *G3*, group 3 (n=14); *G4*, group 4 (n=19); Data normalized via RUV method **(D)** or DeSeq2 median of ratios (MoR) method **(E)** and both analyzed using Mann-Whitney U (\**p* <0.05, \*\**p* <0.01, \*\*\**p* <0.001, \*\*\*\**p* <0.0001). **(F)** Poor prognostic profile demonstrable in high ABCB7-expressing medulloblastoma patients (Cavali *et al.* GSE85217).

**Supplementary Figure 3. ABCB7 repression triggers iron overload. (A and B)** Escalation in both mitochondrial and cytoplasmic free Fe^2+^ demonstrable with miR-1253 expression, rescued by iron chelation with DFO. Images captured at 63X. Confocal images showing calcein AM dye quenching in miR-1253-transfected D425 **(A)** and HDMB03 **(B)** cells indicating high cytosolic labile iron compared to scramble transfected cells. *DFO*, deferoxamine; *NC*, negative control.

**Supplementary Figure 4. ABCB7 repression by MiR-1253 triggers oxidative stress and lipid peroxidation leading to cell death in G3MB cells.** Confocal images showing elevated mitochondrial O_2_ ^•−^ (MitoSOX™ Red, **red**) **(A)** and cytosolic H O (DCFDA, **green**) **(B)** following miR-1253 expression in HDMB03 cells, which could be rescued by ferrostatin. **(C)** A similar trend noted in D425 cells. **(D)** Higher lipid peroxidation (by BODIPY11 staining) in miR-1253 transfected D425 cells. Flow cytometry analysis showing significantly higher O_2_ ^•−^ mediated cell death (representing ferroptosis) in miR-1253-expressing **(E)** D425 and **(F)** HDMB03 cells demonstrable by quantifying cells staining for both Annexin V-Cy5 (apoptosis) and for O_2_ ^•−^ (Mitosox) (Q2). Images captured either at 20X (**A**, **B, C**) or 63X (**D**) magnification. Scale bar 200 µm. Mean ± SD, n=3; *ns*, not significant, \**p*<0.05, \*\**p*<0.01, \*\*\**p*<0.001, \*\*\*\**p*<0.0001. *ERA*, erastin; *FER*, ferrostatin-1; *miR*, miR-1253 overexpression, *NC*, negative control.

**Supplementary Figure 5. GPX4 is a central regulator of ferroptosis that is upregulated in MB. (A)** Schematic showing the important role of GPX4 in mitigating ferroptosis. Image created with <u>Biorender.com</u>. **(B)** Subgroup-specific GPX4 expression assessment by RNA sequencing (log_2_ transcripts per million) of a local medulloblastoma patient cohort (Kanchan *et al.*, GSE148390) showing deregulation in MB tumors, highest in G3MB. *CB*, pediatric cerebellum (n=10); *SHH*, sonic hedgehog (n=6); *G3*, group 3 (n=7); *G4*, group 4 (n=12). **(C)** Subgroup-specific GPX4 expression data from Weishaupt *et al.*, GSE124814; *CB*, normal cerebellum (n=291); *WNT*, wingless (n=118); *SHH*, sonic hedgehog (n=405); *G3*, group 3 (n=233); *G4*, group 4 (n=530); Data normalized via RUV method and analyzed using Mann-Whitney U (\**p* <0.05, \*\**p* <0.01, \*\*\**p* <0.001, \*\*\*\**p* <0.0001). **(D)** Enrichment of key proteins involved in glutathione (GSS, GCLC) and GPX4 (SEPSECS, TRIT1) biosynthesis enriched in G3MB. **(E)** Confocal microscopic images confirming high GPX4 expression in group 3 MB tumors (n=6) compared to pediatric cerebellum (n=6) and colocalization to the mitochondria based on COXIV fluorescence. Images captured at 10X magnification. **(F)** Western blotting analysis showing high *in vitro* GPX4 expression in classic MB cell lines (G3: D341, D425, HDMB03; G3/4: D283) compared to normal human astrocytes (NHA).

**Supplementary Figure 6. ABCB7 repression deplete GSH.** Inhibitory effect of ABCB7 repression by miR-1253 on total glutathione levels in D425 **(A)** and HDMB03 **(B)** and by si-ABCB7 (100 nM) **(C)**. Evaluation of oxidized glutathione (GSH) showing punctuated effects in combination treatment groups (miR + Cis) with rescue in the presence of ferrostatin in miR-1253-transfected HDMB03 cells **(D)** and with ABCB7^KO^ **(E)**.

**Supplementary Figure 7. ABCB7 knockout potentiates cisplatin cytotoxicity in G3MB cells by ferroptosis.** Oxidative stress measured by quantifying **(A)** mitochondrial O_2_ ^•−^ (MitoSOX™ Red, **red**) and **(B)** cytosolic H_2_O_2_ (DCFDA, **green**) in ABCB7^KO^ HDMB03 cells showing potentiating effect of cisplatin, and inhibited by ferrostatin-1. Images captured at 20X magnification. Scale bar 200 µm. **(C)** As measured by Image-iT® Lipid Peroxidation Kit, confocal images showing the highest lipid peroxidation in combination treatment groups (ABCB7^KO^ + Cis), again rescued by ferrostatin-1, in ABCB7^KO^ HDMB03 cells. Images captured at 63X magnification. Mean ± SD, n=3; \**p* <0.05, \*\**p* <0.01, \*\*\**p* <0.001, \*\*\*\**p* <0.0001.

