## Supplementary figures and images for "Repressing ABCB7 Potentiates Cisplatin Response in Pediatric Group 3 Medulloblastoma by Triggering Ferroptosis"

### Supplementary Figure 1

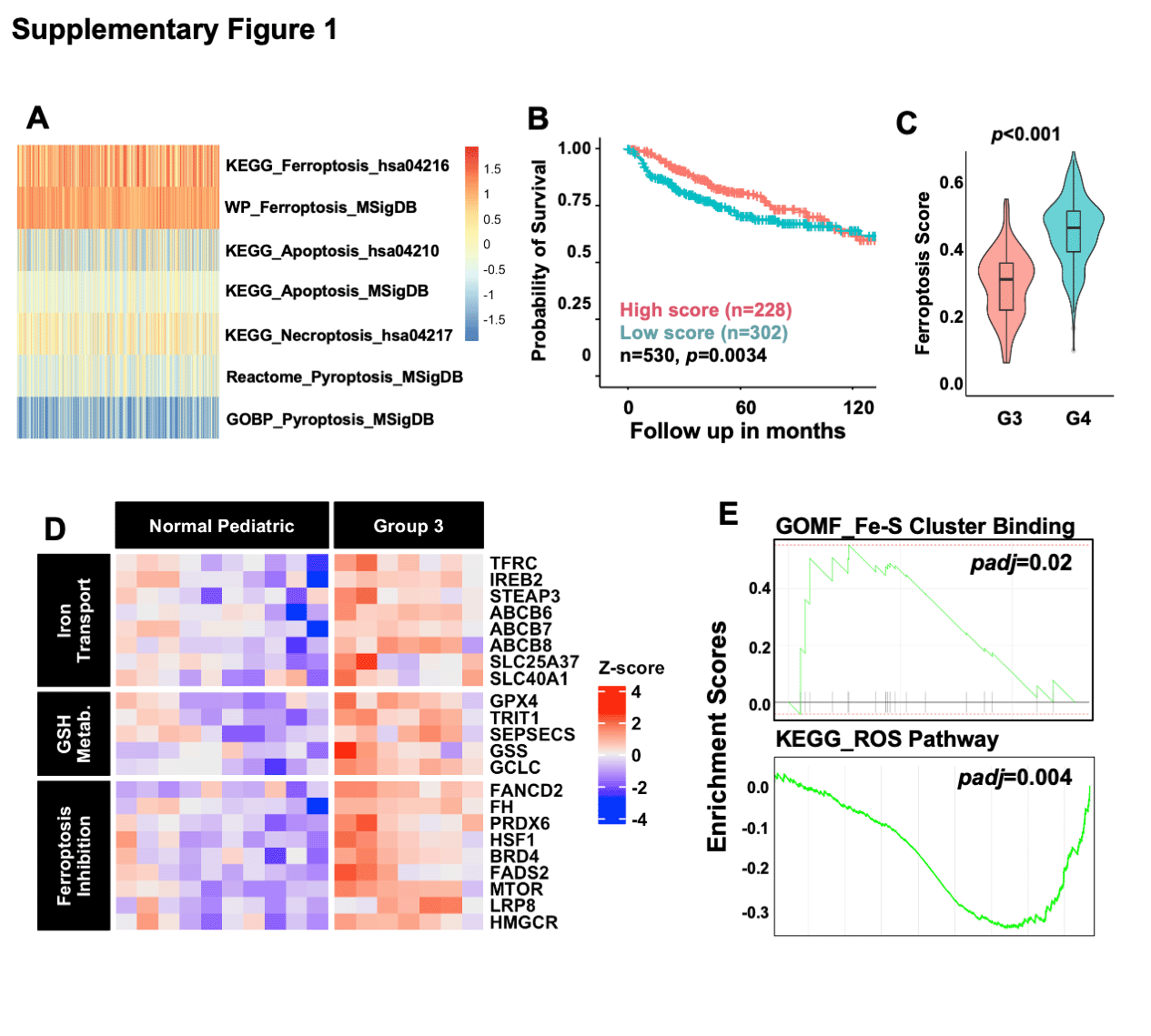

### Supplementary Figure 2

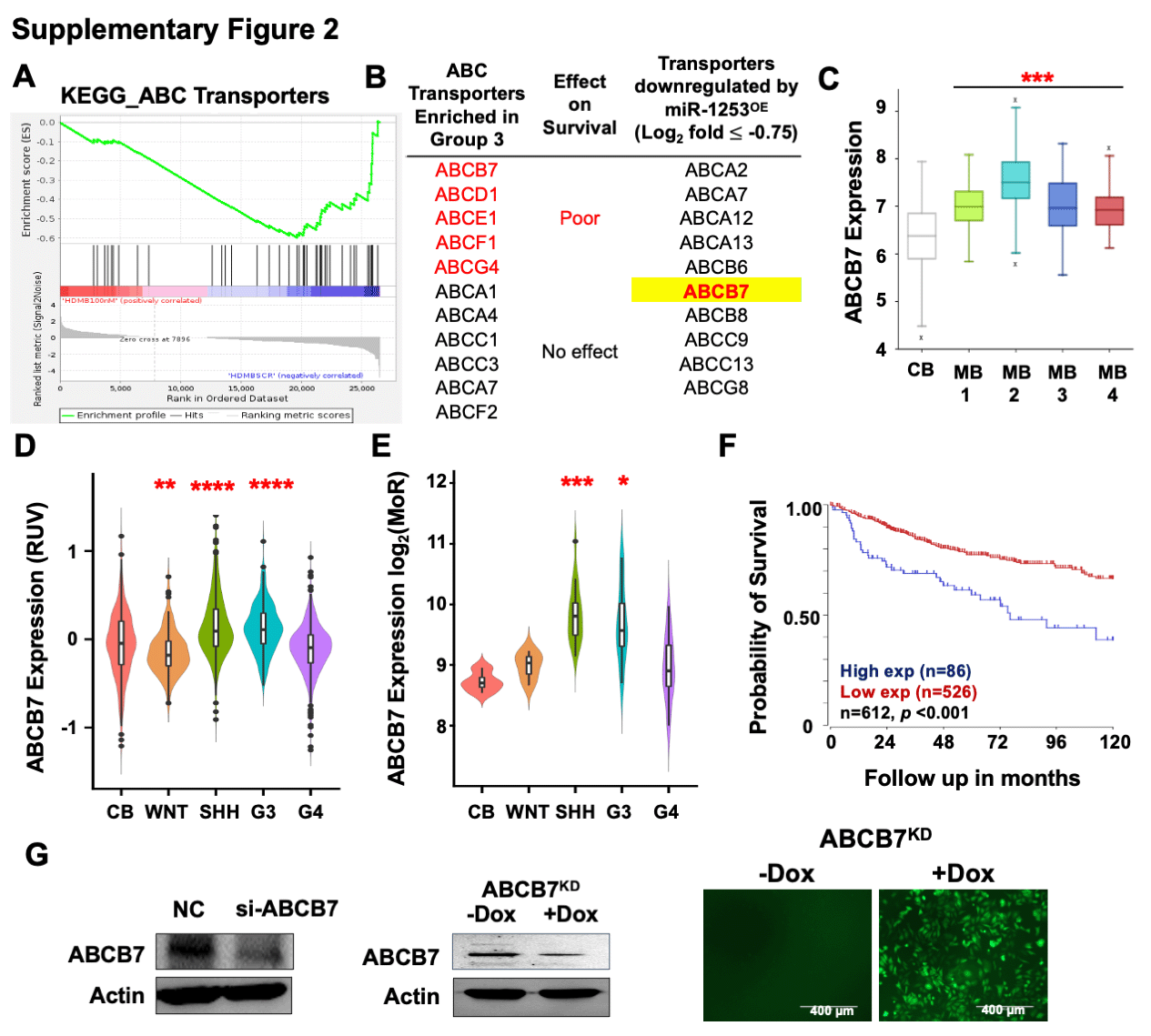

### Supplementary Figure 3

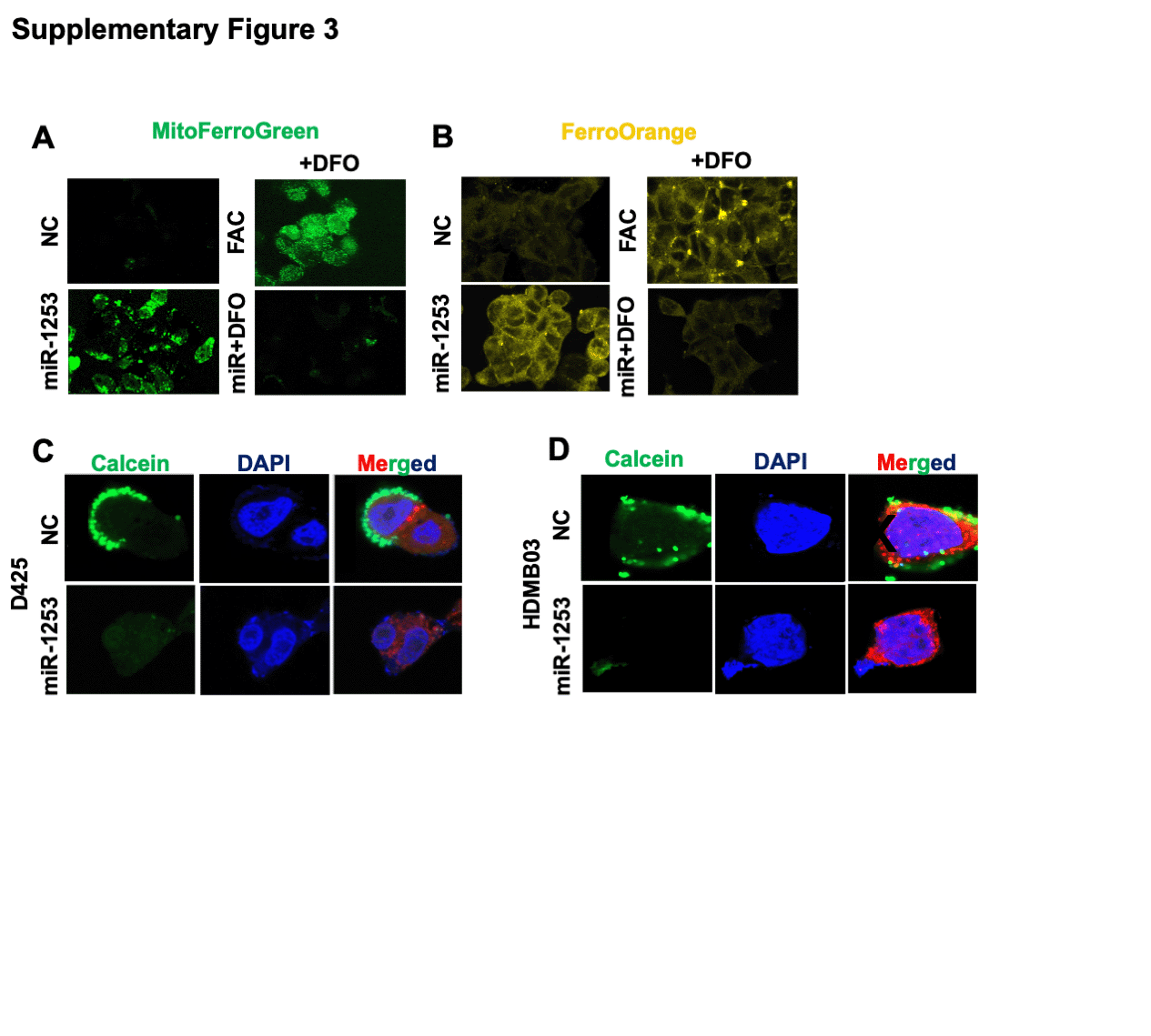

### Supplementary Figure 4

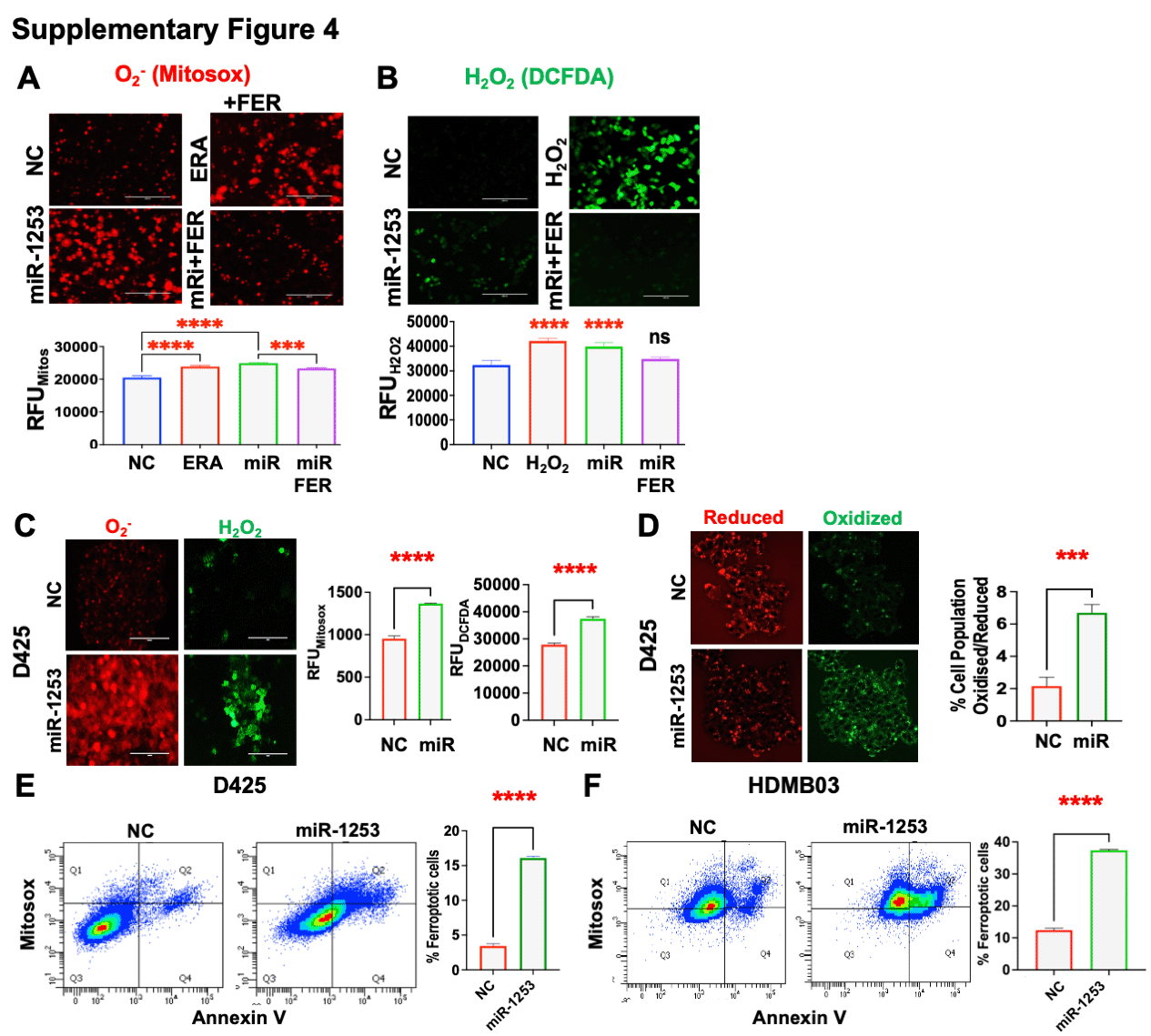

### Supplementary Figure 5

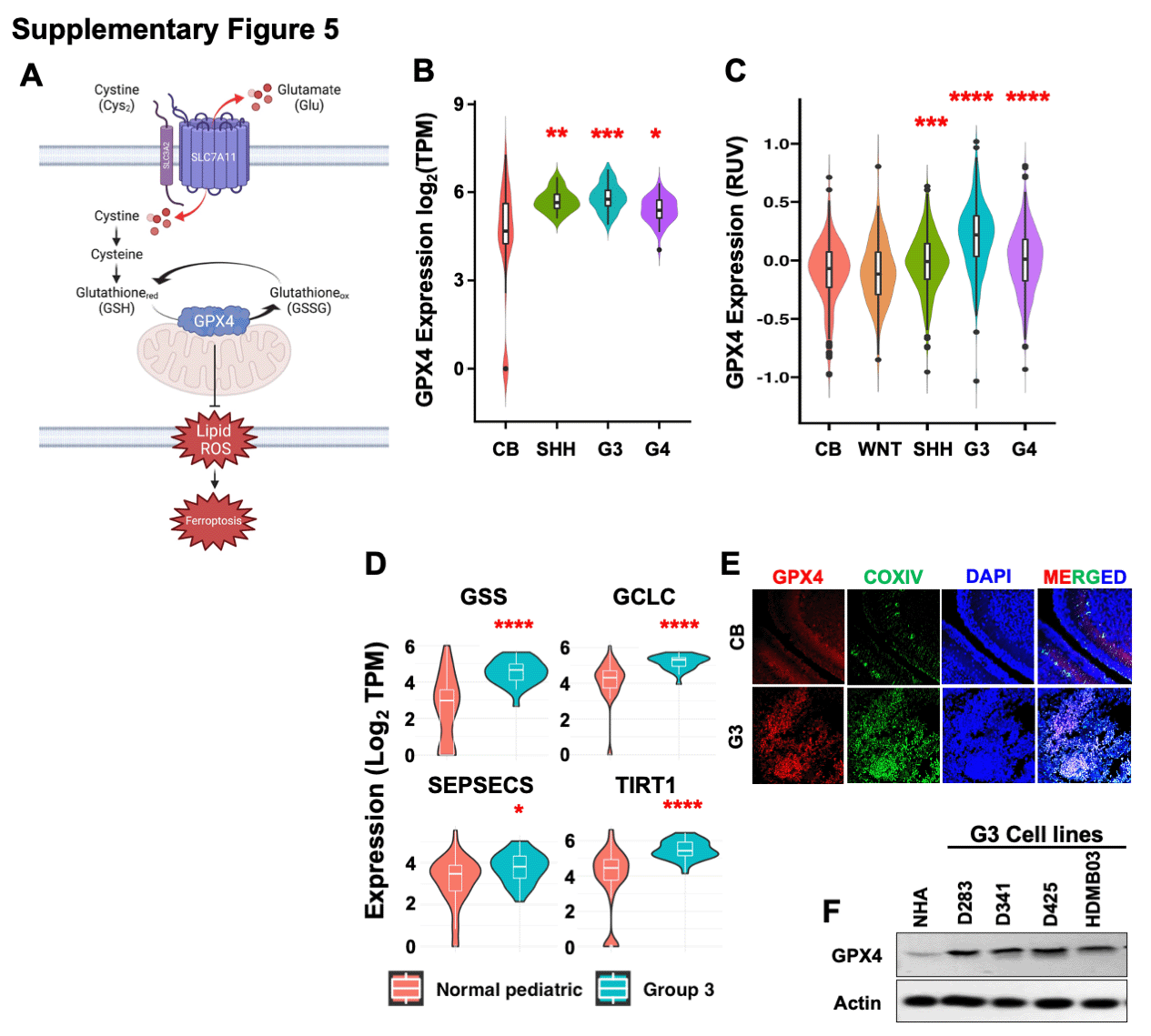

### Supplementary Figure 6

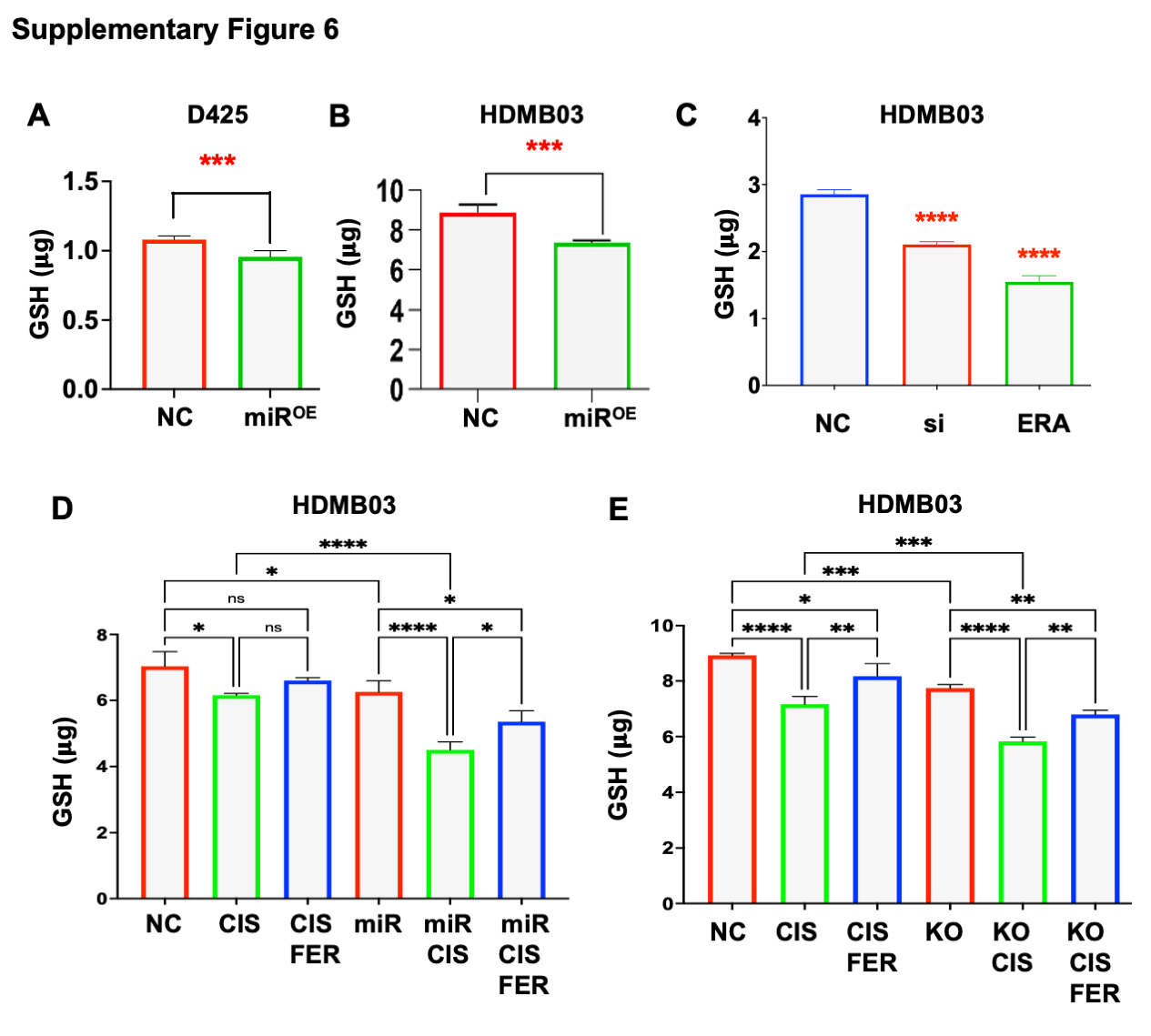

### Supplementary Figure 7

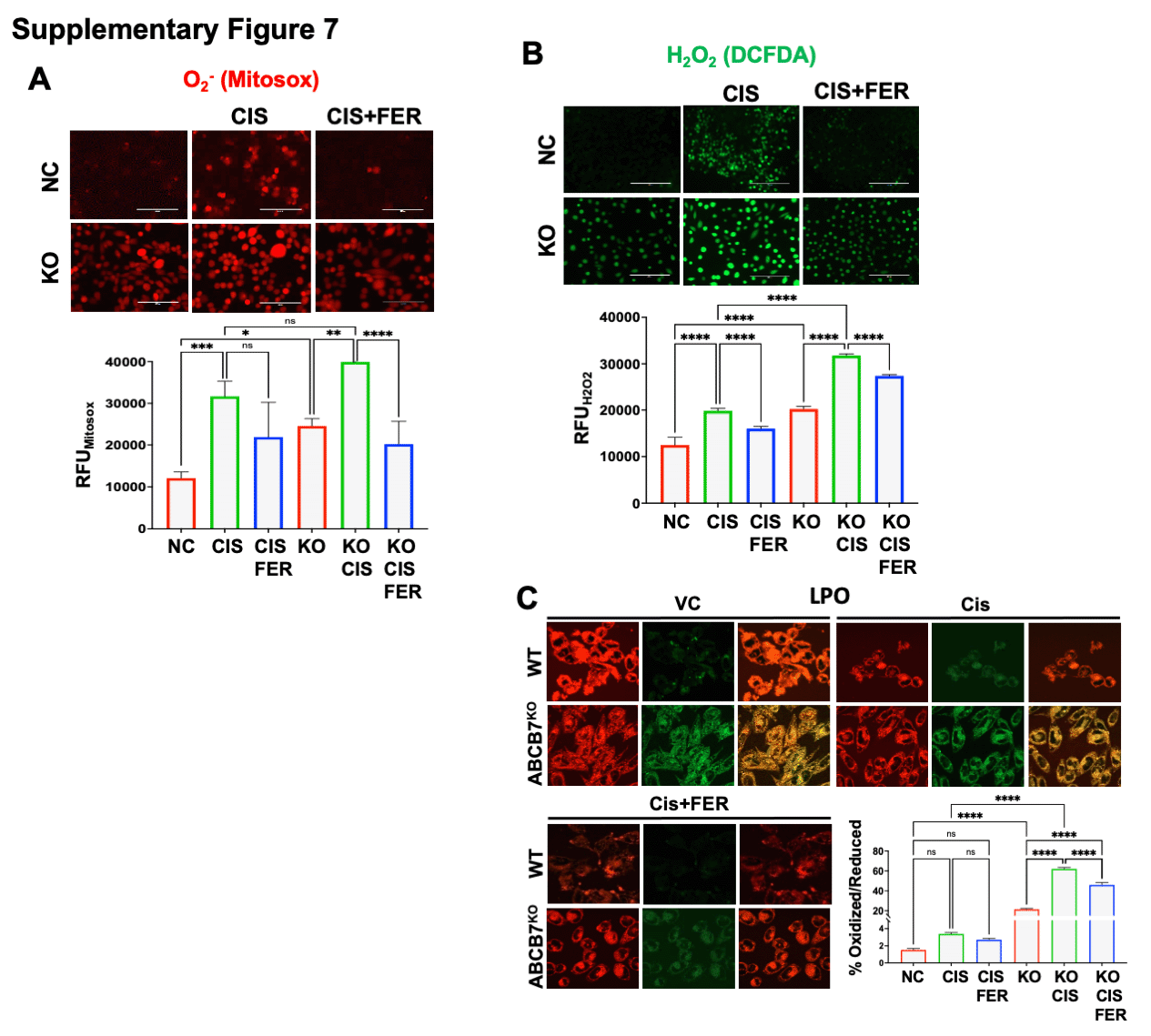
